# Xylazine is an agonist at kappa opioid receptors and exhibits sex-specific responses to naloxone administration

**DOI:** 10.1101/2023.09.08.556914

**Authors:** Madigan L. Bedard, Jackson G. Murray, Xi-Ping Huang, Alexandra C. Nowlan, Sara Y. Conley, Sarah E. Mott, Samuel J. Loyack, Calista A. Cline, Caroline G. Clodfelter, Nabarun Dasgupta, Bryan L. Roth, Zoe A. McElligott

## Abstract

Xylazine has been found in the unregulated drug supply at increasing rates, usually in combination with fentanyl. It has become critical to understand its basic pharmacology, how it impacts behavior, and how it interacts with fentanyl in rodent models of opioid administration. Despite commentary from scientists, politicians, and public health officials, it is not known if xylazine impacts the efficacy of naloxone, the opioid receptor antagonist used to reverse opioid induced respiratory depression. Furthermore, few studies have examined the effects of xylazine alone, without co-administration of ketamine. Here, we examine the impact of xylazine alone and in combination with fentanyl on several key behaviors in male and female mice. We demonstrate differential locomotor responses by dose and sex to xylazine. Surprisingly, our results further indicate that naloxone precipitates withdrawal from xylazine and a fentanyl/xylazine combination, in both sexes, with enhanced sensitivity in females. Further, we show that xylazine is a full agonist at the kappa opioid receptor, a potential mechanism for its naloxone sensitivity.

**One-Sentence Summary:** We present surprising new insights into xylazine and fentanyl pharmacology with immediate implications for clinical practice and frontline public health.

## INTRODUCTION

Human exposure to the veterinary anesthetic xylazine has been reported intermittently in Spain (*1*), Germany (*2*), Canada (*3*), and the United States (*4, 5*) since the 1970s, often associated with occupational exposure and intentional self-harm. Sustained use of the liquid veterinary formulation for euphoric effect was documented in Puerto Rico starting around 2001 (*6*), with sporadic detection in seized street drugs first on the east coast of the United States mainland from 2006 onwards, and in California for at least the last 4 years (*7, 8*). The complex interplay between illicitly manufactured xylazine, heroin, fentanyl, and methamphetamine supply can be traced to power shifts among drug trafficking organizations, exacerbated by international drug control policies; overseas chemical manufacturers have responded to demand for fentanyl alternatives driven by consumer dissatisfaction with the potent opioid (*1, 6, 7, 9, 10*). Currently, xylazine is predominantly found in powder forms of unregulated street drugs in many (but not all) regions of the United States, and mostly (but not exclusively) with illicitly manufactured fentanyl (*1, 6, 7*). In recognition, the federal government designated “fentanyl adulterated or associated with xylazine” as an emerging drug threat in April 2023 (*6, 7*). However, xylazine and fentanyl co-exposure is not an exclusively American phenomenon: the earliest documented xylazine-fentanyl co-ingestion was accidental in a farmworker in New Zealand in 1984 (*11*), and a xylazine-fentanyl-heroin overdose death was reported in the United Kingdom in 2023 (*12*). Clinical management of human xylazine exposure is made difficult by disfiguring and lingering skin ulcers, a distinctive agitated withdrawal syndrome, and lack of approved antidote or withdrawal support medications (*13, 14*). Ultimately, the limited pharmacological understanding of xylazine, in conjunction with the lack of an approved antidote, has hampered effective responses to this emerging threat.

Despite having similar sedative effects, fentanyl and xylazine previously have been thought to act on different G protein-coupled receptors (GPCRs). Canonically, xylazine acts on the alpha-2-adrenergic receptor (α_2_-AR) whereas fentanyl acts on mu, kappa, and delta opioid receptors (μOR, κOR, δOR respectively). As xylazine has been increasingly found in the unregulated drug supply, there have been reports of worsened overdoses attributed to mixtures of fentanyl and xylazine (*15–17*). A general assumption has been that due to the presence of xylazine, these overdoses are not responsive to naloxone (*18, 19*), an opioid receptor antagonist used to alleviate respiratory depression induced by opioids. Some evidence has indicated, however, that xylazine may also act on other receptors (*20*), though it has not been directly tested *in vitro* nor *in vivo* until now.

Much of the preclinical veterinary research has focused on xylazine’s sedative effects in combination with ketamine (*21, 22*), and few studies have investigated xylazine on its own or in the context of reward learning (*23, 24*). Additionally, these studies did not report locomotor effects to control for potential sedative effects of α_2_-AR agonists which could potentially impede learning mechanisms in rodent models. Recently, Khatri et al. found that xylazine depressed fentanyl self-administration in male and female rats (*25*). However, α_2_-AR agonists (e.g. clonidine), despite being commonly used to treat opioid withdrawal, may have dependence potential themselves (*26–34*). Here, we sought to better understand the effects of xylazine alone and determine if it alters the fentanyl-withdrawal experience in both male and female C57BL/6J mice.

## METHODS

### Subjects

All procedures described were approved by the University of North Carolina at Chapel Hill Institutional Animal Care and Use Committee (IACUC). Male and female C57BL/6J mice aged 8-12 weeks were purchased from The Jackson Laboratory and maintained on a normal 12-hour light-dark cycle, receiving food and water *ad libitum*. All behavioral experiments were conducted during the light cycle.

### Drugs

All drugs were delivered in 0.9% sterile saline solution and injected either subcutaneously (SC) or intraperitoneally (IP) based on body weight. Fentanyl citrate was purchased from Spectrum Chemical MFG Corp (Gardena, California) and delivered at 0.1 mg/kg IP. Naloxone hydrochloride dihydrate was purchased from Sigma-Aldrich (St. Louis, MO) was delivered at 1 mg/kg SC. Xylazine injectable solution (20 mg/mL) was purchased from Covetrus, Inc. (Portland, ME) and used at either 0.5, 1, or 3 mg/kg IP. Atipamezole hydrochloride (Tocris, Bristol, UK) was used at 1 mg/kg SC. Experimenters were blind to the administered doses whenever possible.

### Acute Locomotion

Mice (N=10-12/group/sex) were brought to the experimental room 30 minutes prior to testing to acclimate. They were weighed and administered one of four possible doses of xylazine (0, 0.5, 1, or 3 mg/kg) via IP injection. Locomotor activity was monitored for 1 hour immediately following injection using the Accuscan Fusion SuperFlex Open Field system (https://omnitech-usa.com/product/superflex-open-field/). The animals were allowed to freely explore the 16×16-inch acrylic arena and their movement was tracked by their interference of infrared beams.

### Precipitated Withdrawal

Mice (N=10/group/sex) were injected in their home cage room with fentanyl citrate (0.1 mg/kg, IP), xylazine injectable solution (0.5 mg/kg IP), fentanyl citrate + xylazine injectable solution (0.1 mg/kg + 0.5 mg/kg IP), or sterile saline (0.9%, equivolume IP). Mice were then placed back on their colony rack. 2 hours later, mice were moved to a procedure room where withdrawal was precipitated with naloxone hydrochloride (1 mg/kg SC) or atipamezole hydrochloride (1 mg/kg SC) and withdrawal behaviors were scored for 10 minutes following injection. The following withdrawal behaviors were scored: escape jumps, paw tremors, jaw tremors, wet dog shakes, fecal boli count, grooming, and abnormal posture. Mice were returned to their home cages and precipitated withdrawal was repeated for 2 more days (*35–37*). Total counts for each behaviour were recorded and a z-score was computed for each animal on each day of withdrawal (*38*). On the third day of withdrawal, mice were perfused for immunohistochemistry 75-90 minutes following antagonist administration.

### Immunohistochemistry and Microscopy

75-90 minutes after naloxone administration, mice were perfused with 4% paraformaldehyde (in 0.01 M PBS). Brains (N=4-5/group/sex) were removed and remained in fixative for 24 h followed by 30% sucrose (in 0.01 M PBS) for at least 48 h before being sectioned by cryostat (CM 3050S). Sections were incubated overnight at 4°C in blocking solution containing primary antibody: rabbit anti-c-Fos (1:3000; Synaptic Systems). The following day, sections were incubated in HRP-conjugated goat anti-rabbit IgG (1:200; Perkin Elmer) for 30 minutes then incubated in Cy3 diluted in TSA amplification diluent (1:50) for 10 minutes. Sections were then incubated in DAPI diluted in PBS (1:1000) as a counterstain. Images were collected and processed on an Evident Sci VS200 Slide Scanner at 10x magnification. QuPath was used for cell counting and analysis.

### Radioligand binding and GPCR functional assays

Radioligand binding assays and functional GPCR assays (PRESTO-Tango and GloSensor) were conducted by the Psychoactive Drug Screening Program (<u>PDSP</u>) as described in Besnard et al. (2012), Kroeze et al. (2015), Salas-Estrada et al. (2023), and Olsen et al. (2020) (*39–42*).

### Statistics

All statistics were calculated using GraphPad Prism version 10.0.0. Data is represented as mean ± SEM unless otherwise noted in the figure legend. Indications of statistical significance followed GraphPad p-value style (p > 0.05 ns, p ≤ 0.05 *, p ≤ 0.01 **, p ≤ 0.001 ***, p ≤ 0.0001 ****). Figure legends include information on the specific tests run.

## RESULTS

### Identification of non-sedative doses of xylazine

Few studies in mice have investigated the sedative effects of xylazine administered alone (i.e., without the addition of ketamine or other anesthetics) (*43*). Because sedation alters locomotor activity, it is necessary to determine a non-sedative dose to use in future behavioral experiments. Typically, 10 mg/kg dose of xylazine is used with ketamine for anesthesia (*21, 22*). Previous behavioral studies have tested doses as low as 2.5-3 mg/kg in mice but did not validate that this was a non-sedative dose (*43, 44*), so we probed a lower range of 0, 0.5, 1, and 3 mg/kg xylazine in the open field assay. We found that 3 mg/kg xylazine resulted in decreased distance traveled in both males and females compared to saline and 0.5 mg/kg, and in females compared to 1 mg/kg (Fig. 1A). A dose of 3 mg/kg also decreased the % ambulatory time compared to the other three doses in both males and females (Fig. 1B). In males but not females, the 1 mg/kg dose decreased % time ambulating compared to saline (Fig. 1B). Additionally, none of the doses resulted in a reduction in the average velocity for either sex across the full 60-minute trial. However, at the lowest dose (0.5 mg/kg), females exhibited significantly higher average velocity than the males in the second 30 minutes of the trial (Fig. 1F).

**Fig. 1.**
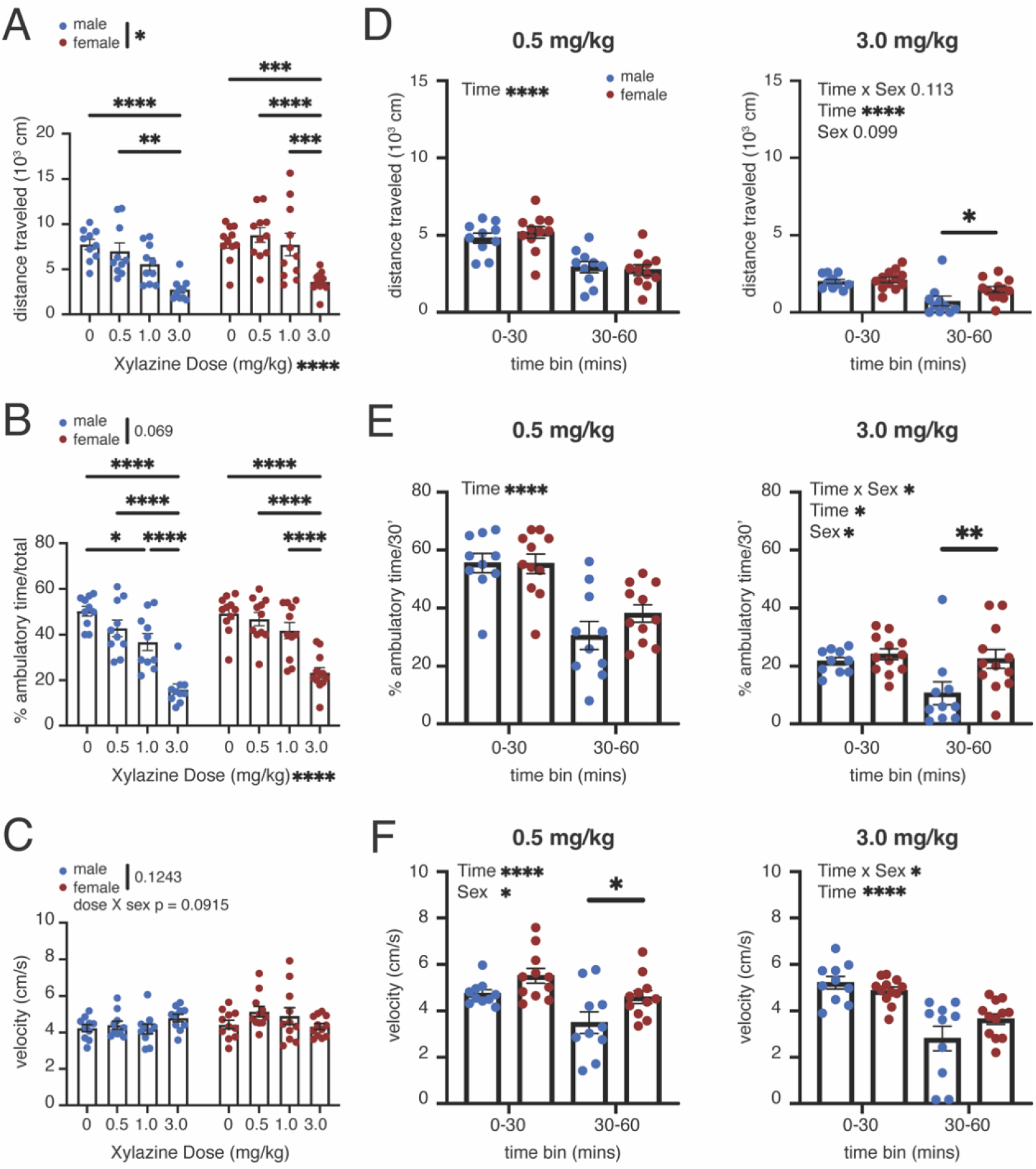
Effect of acute xylazine administration on locomotor activity. (A) Cumulative distance traveled, (B) % ambulatory time, and (C) average velocity of male and female mice administered saline or xylazine (0.5, 1.0, or 3 mg/kg). (D) Distance traveled, (E) % ambulatory time, and (F) velocity split into 30-minute bins for 0.5 mg/kg (left panel) and 3.0 mg/kg (right panel). (A-C) 2-way ANOVAs (Dose X Sex) with Tukey’s post-hoc. (D-F) 2-way ANOVAs (Time x Sex) with Sidak’s post-hoc.

Xylazine’s onset of action is estimated to be about 10-15 minutes and exploratory behavior naturally declines over time due to intrasession habituation. To examine the temporal effects of xylazine on locomotor activity, we further analyzed the data in both 10-minute (Fig. S1) and 30-minute time bins (Fig. 1D-F). As expected, distance traveled, % time ambulatory, and velocity generally declined across time for both male and female mice at all doses (Fig. S1A-B). However, we observed interesting sex differences that emerged across the time course of the experiment. Surprisingly, at low doses of xylazine (0.5 mg/kg - 1 mg/kg) females traveled a greater distance and at a higher average velocity compared to their male counterparts (Fig. 1F). At 3 mg/kg, female velocity dropped precipitously in the first half of the trial, but recovered such that distance traveled and ambulatory time significantly exceeded the males in the second half of the trial (Fig. 1D-E). These data confirm that xylazine can exert sedative effects at doses as low as 1 mg/kg in male mice and 3 mg/kg in female mice. Our results further suggest that at 3mg/kg there are significant sex differences in the time course of and recovery from the sedative effects of xylazine. We chose to proceed with 0.5 mg/kg xylazine because it was non-sedative in all measures of both sexes and even slightly hyperlocomotive in female mice (Fig. 1).

### Naloxone- and Atipamezole-Precipitated Withdrawal

Withdrawal from reinforcing substances is a critical component of the addiction cycle (*45*– *47*). Previously, we and others have used a 3-day repeated precipitated morphine withdrawal model to demonstrate that somatic symptoms exacerbate across withdrawal sessions (*35–37, 48–51*). Furthermore, we have shown that this model results in sleep disturbances and promotes long-lasting sex-dependent behavioral adaptations in both male and female mice over six weeks into forced abstinence (*35*). Here we adapted our model to fentanyl withdrawal and explored if fentanyl/xylazine co-administration would impact the development of the withdrawal syndrome. Male and female mice were administered (IP) either saline (equal volume), fentanyl (0.1 mg/kg), xylazine (0.5 mg/kg), or a combination of fentanyl/xylazine (0.1 and 0.5 mg/kg respectively). Two hours later, mice received an injection of either naloxone (1 mg/kg SC) or atipamezole (1 mg/kg SC, the α_2_-AR antagonist used by veterinarians to reverse xylazine anesthesia; Fig. 2).

**Fig. 2.**
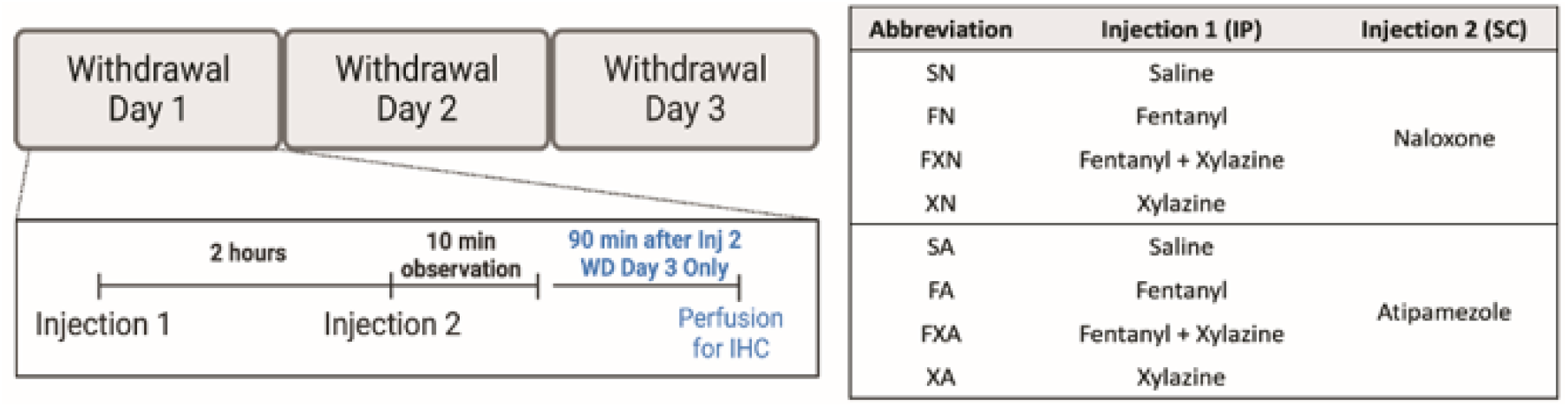
Precipitated Withdrawal Paradigm and Treatment Groups.

In contrast to the conventional concept that ‘xylazine is not affected by naloxone’, we found that females that had only received xylazine demonstrated significant global somatic withdrawal scores (shown as z-scores (*38*)) following naloxone administration, which sensitized over three days (Fig. 3A,B). Indeed, across the 3-day paradigm, xylazine withdrawal was of equal severity to fentanyl withdrawal in females (Fig. 3A). Furthermore, female mice showed the most dramatic somatic withdrawal when fentanyl and xylazine were combined. In contrast, male mice demonstrated the most dramatic withdrawal to fentanyl alone, and the fentanyl/xylazine combination did not alter the degree of withdrawal experienced (Fig. 3A,B). We also considered that the sexes and treatment groups might experience different types of somatic withdrawal symptoms. To assess this, we plotted the average z-score for each individual behavior on withdrawal day 3 (Fig. 3B,D). Interestingly, the most severe symptom for both sexes was paw tremors in the fentanyl/xylazine combination group. Regardless, females exhibited multiple withdrawal behaviors that were enhanced in the fentanyl/xylazine combination compared to either the fentanyl or xylazine groups. In both sexes, the addition of xylazine decreased the fecal bolus count relative to the fentanyl group (Fig. 3B). Males displayed a more robust increase in escape jumps due to the fentanyl/xylazine combination than females, but females saw increases in wet dog shakes and abnormal posture between fentanyl/xylazine and fentanyl (Fig. 3B).

**Fig. 3.**
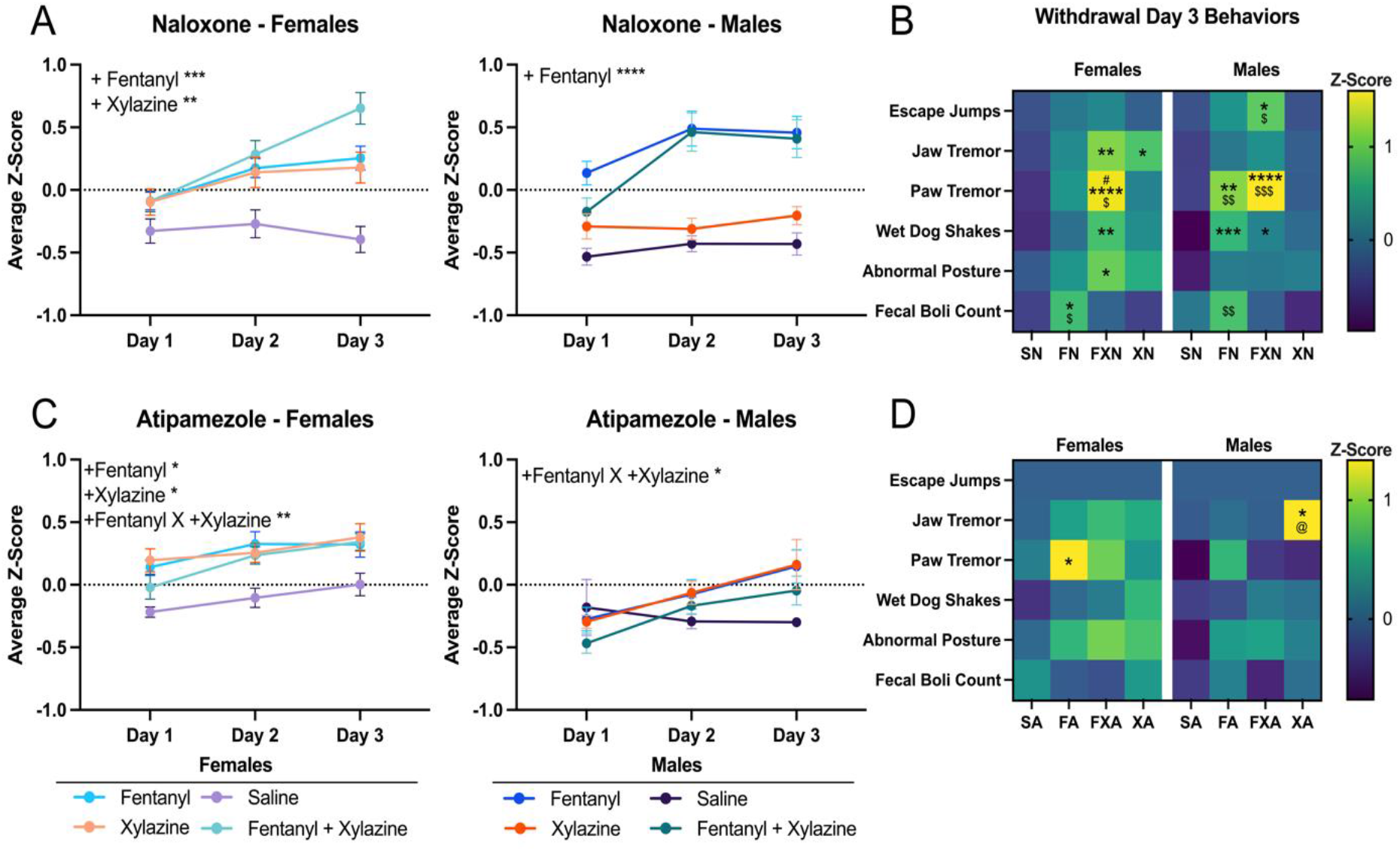
Effect of acute xylazine administration on locomotor activity. Global scores are shown as average z-score ± SEM. (A) Average z-scores of female and male mice over three days of naloxone-precipitated withdrawal. (B) Heatmap of average z-scores on day three of withdrawal for individual behaviors. (C) Average z-scores of female and male mice over three days of atipamezole-precipitated withdrawal. (D) Heatmap of average z-scores on day three of atipamezole-precipitated withdrawal for individual behaviors. (A & C) 3-way ANOVAs (Day X Addition of Fentanyl X Addition of Xylazine) P=0.05*, 0.01**, 0.001***, 0.0001****. Main effects p-values and Tukey’s post-hoc shown in Fig. S2. (B & D) 2-way ANOVAs (Tx Group X Behavior) with Tukey’s post-hoc, P=0.05 where * (vs. saline), # (vs. fentanyl), @ (vs. fentanyl/xylazine), and $ (vs. xylazine). SN=saline-naloxone; FN=fentanyl-naloxone; FXN=fentanyl/xylazine-naloxone; XN=xylazine-naloxone; SA=saline-atipamezole; FA=fentanyl-atipamezole; FXA=fentanyl/xylazine-atipamezole; XA=xylazine-atipamezole.

Similarly to the results with naloxone, we were surprised to observe that atipamezole was able to induce precipitated withdrawal behaviors from animals exposed to fentanyl alone. Females exhibited similar levels of withdrawal in fentanyl, xylazine, and fentanyl/xylazine groups in response to atipamezole (Fig. 3C). Males showed lesser degrees of atipamezole-induced withdrawal overall compared to females. Interestingly males responded similarly to xylazine and fentanyl, but the combination of the two dampened the effects. Finally, males showed no withdrawal symptom sensitization to saline-atipamezole over the three days, but the females did (Fig. 3C).

These data indicate a hyposensitivity of males to xylazine compared to females at this low dose and that female responses to fentanyl withdrawal can be enhanced by the addition of xylazine. Further, females exhibited increased sensitivity to the α_2_-AR antagonist than males did, indicating a sex difference in adrenergic systems.

### Male and female mice exhibit differential c-Fos expression following naloxone-precipitated withdrawal

75-90 minutes following naloxone administration on the final day of withdrawal, mice were perfused for immunohistochemistry. The immediate early gene, c-Fos, is expressed by active neurons, reaching peak expression approximately 90 minutes after activity (*52, 53*). Within the regions analyzed, significant differences between treatment groups were observed in the locus coeruleus (LC), dorsal bed nucleus of the stria terminalis (dBNST), and lateral central nucleus of the amygdala (lCeA) in females and in the LC and basolateral amygdala (BLA) in males (Fig. 4).

**Fig. 4.**
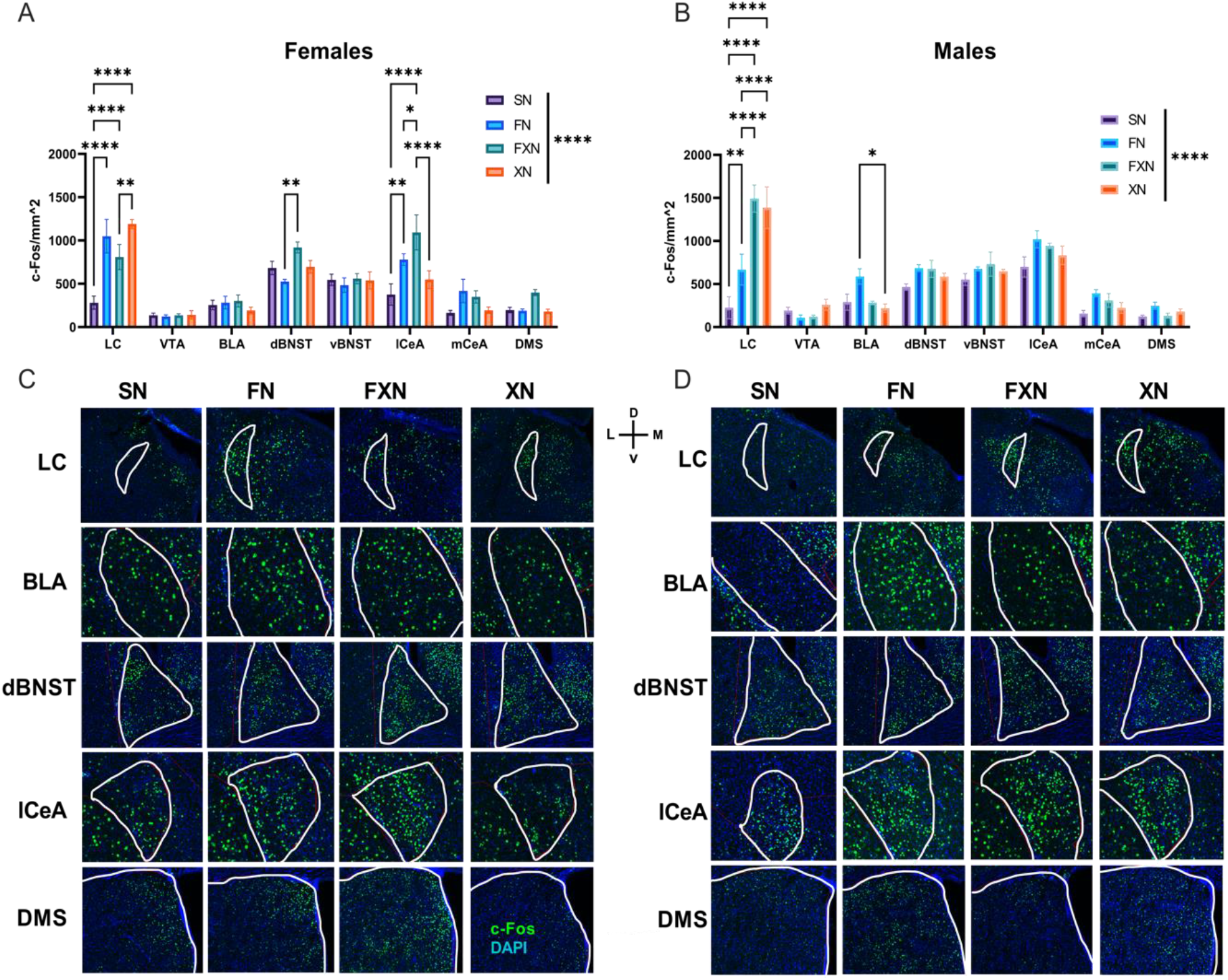
Quantification of c-Fos expression following precipitated withdrawal. (A) Female c-Fos expression displayed as number of positive cells per mm^2. (B) Male c-Fos expression displayed as number of positive cells per mm^2. (C) Representative images of female regions of interest. (D) Representative images of male regions of interests. (A & B) 2-way ANOVAs (Tx group X ROI) with Tukey’s post-hoc. P=0.05*, 0.01**, 0.001***, 0.0001****.

LC regional c-Fos expression was significantly higher in female mice that received fentanyl alone and xylazine alone compared to female mice that received the fentanyl/xylazine combination (Fig. 4A). Intriguingly, male mice exhibited a different LC c-Fos expression response wherein the male mice that received xylazine alone and the fentanyl/xylazine combination had significantly higher c-Fos expression than the male mice that received only fentanyl (Fig. 4B). In both sexes, the three treatment groups exhibited significantly higher c-Fos expression than the mice that received saline (Fig. 4A,B).

Female mice displayed significant c-Fos expression differences in a few additional regions of interest. In the dBNST, the female mice that received the fentanyl/xylazine combination had significantly higher c-Fos expression as compared to the female mice that received fentanyl alone. Interestingly, no differences were observed between the saline mice and those that received xylazine, suggesting a dBNST effect that is driven primarily by fentanyl administration (Fig. 4A). Differences in c-Fos expression within the lCeA were also observed in female mice, wherein the fentanyl/xylazine combination displayed significantly greater c-Fos expression than all other treatment groups. Mice that received fentanyl alone had greater expression than the saline mice, and no differences were observed between the saline mice and those that received xylazine alone (Fig. 4A).

In males, only one additional region showed differences in c-Fos expression. In the BLA, males that received fentanyl alone displayed significantly greater c-Fos expression than those that received xylazine alone, while the addition of xylazine to the fentanyl in the combination group reduced c-Fos expression to saline levels (Fig. 4B).

### Characterization of xylazine binding and activity profiles

Xylazine is canonically believed to be an α_2_-AR agonist, though its activity at different receptors has not been tested. To begin to screen, we tested xylazine (10 μM) across a host of common drug targets for radioligand binding activities. Xylazine binds α_2_-ARs, as well as serotonin 7A receptor (5-HT_7A_R), dopamine D1 (D1R), kappa opioid receptor (κOR), sigma 1 receptor (σ1R), and sigma 2 receptors (σ2R) (Fig. S3A). We also tested xylazine in the PRESTO-tango GPCRome screen for potential agonist activity at 320+ human GPCRs. These data indicated xylazine (10 μM) activates α_2A_-ARs as expected and binds several receptors including: α_2B_-AR, α_2C_-AR, κOR, and dopamine D2 receptor (Fig. 5D). The results at the κOR were validated in additional assays. A Gi-GloSensor assay showed that xylazine acts as a full agonist at the κOR with a potency of 1.4 μM (pEC_50_ of 5.86) and was as efficacious (although less potent) as the naturally occurring κOR agonist, salvinorin A (Fig. 5H). Xylazine was also able to displace a radiolabeled κOR agonist ^3^H-U69593 (Fig. G, pK_i_ = 6.33 ± 0.02, K_i_ = 0.47 μM). Xylazine only showed activity at the κOR, not the μ or δ opioid receptors as shown in concentration response curves (Fig. 5A-C).

**Fig 5.**
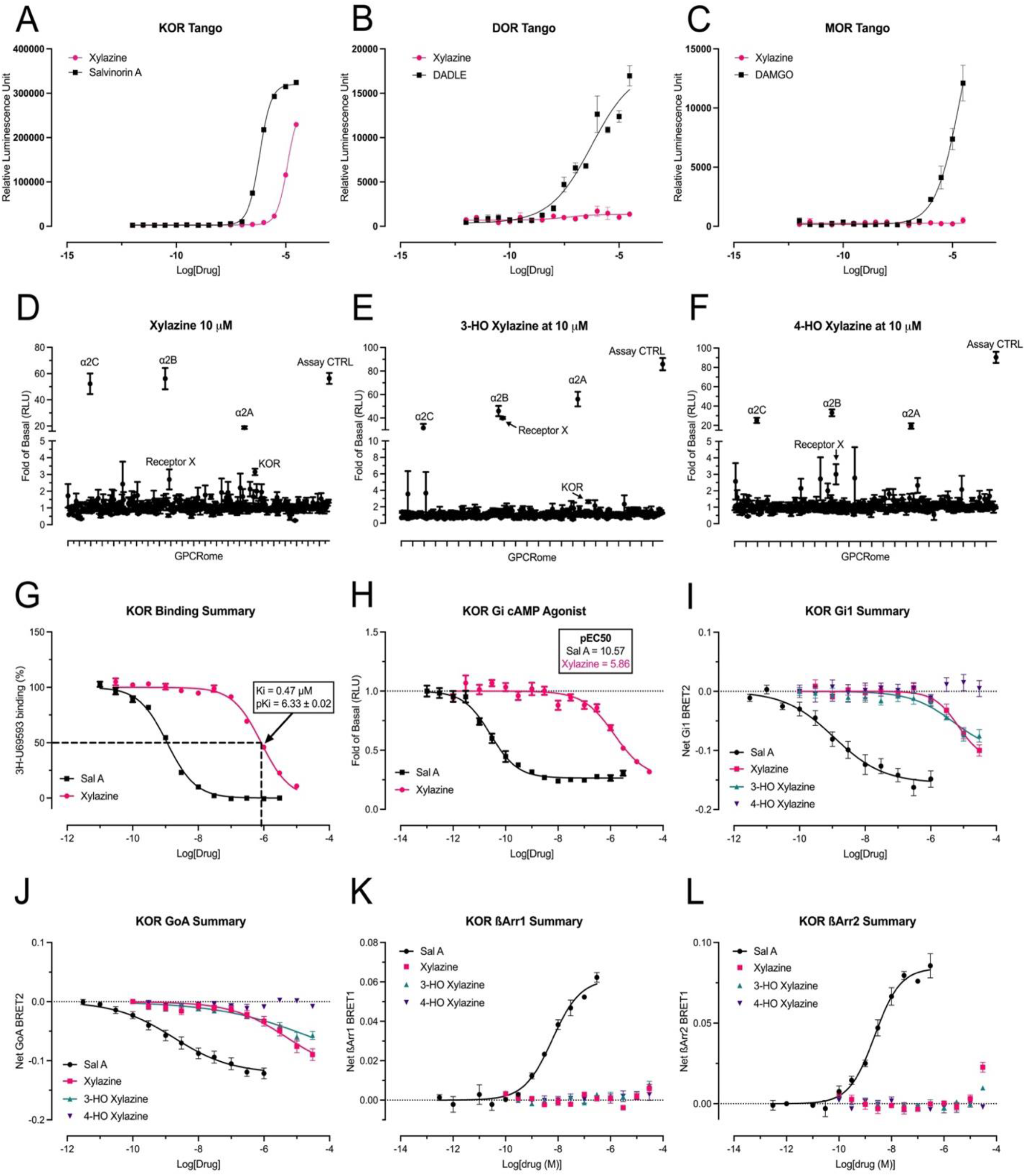
Xylazine acts as an agonist at the kappa opioid receptor. (A-C) Concentration response assays for xylazine activity at κOR (A), δOR (B), and μOR (C) all shown with known reference agonists. (D-F) Fold of basal luminescence of 10 μM xylazine (D) and its primary metabolites (E: 3-hydroxy-xylazine; F: 4-hydroxy-xylazine) tested in the PRESTO-Tango assay against 300+ human GPCRs, potential hits are labelled. (G) Radioligand competitive binding assay confirms kappa receptor binding. (H) Xylazine shows full agonism at κOR in Gi-GloSensor assay at higher concentrations than the positive control Salvinorin A. (I-L) Compounds tested in TPRUPATH assay for activities in either G proteins or ß-arrestin pathways.

The major metabolites of xylazine, 3-hydroxy- and 4-hydroxy-xylazine (*54, 55*), were also tested for activity at μ, δ, and κ (Fig. S3B-E). Xylazine and both metabolites showed Gi agonist activity at kappa but not the other opioid receptors in these assays. 3-hydroxy-xylazine was as efficacious, though less potent, as xylazine and salvinorin A. 4-hydroxy-xylazine was less efficacious at kappa overall. Bioluminescence Resonance Energy Transfer 2 (BRET2) assays (TRUPATH (*42*)) were used to identify potential bias activity among inhibitory G proteins (Gi1, GoA and Gz) and arrestin signaling pathways by xylazine and the metabolites. Xylazine and 3-hydroxy-xylazine showed similar activation of Gi1 and GoA pathways while 4-hydroxy showed no activity (Fig. 5L,J). Interestingly, xylazine and the metabolites showed no activity in the ß-arrestin 1 pathway, while xylazine and 3-hydroxy-xylazine showed weak activity at 30 μM point in the ß-arrestin 2 pathway, consistent with above Tango results, which measures ß-arrestin 2 recruitment mediated luciferase reporter activity (Fig. 5K,L). These findings indicate xylazine and 3-hydroxy-xylazine are both G protein biased agonists at κOR.

## DISCUSSION

Cycles of drug exposure and withdrawal are thought to be critical to the development of substance use disorders (*46, 47*). The dramatic increase of xylazine in the drug supply in recent years prompts the need to understand how xylazine may interact both alone and in conjunction with fentanyl to alter behavioral and physiological responses. Here, we report the first xylazine dose-response locomotor study in male and female mice as well as the first assessment of adrenergic- and opioid-receptor antagonist-precipitated withdrawal symptoms following, xylazine, fentanyl, and xylazine/fentanyl administration in mice. These experiments show that males and females are differentially sensitive to xylazine. We found that female mice were less sensitive to the motor-suppressing effects of xylazine contrary to the recent findings in rats reported by Khatri et al. (2023), potentially due to their use of repeated dosing of xylazine or species differences (*25*). Using a modified version of our 3-day precipitated withdrawal model (*35–37*), we show xylazine is indeed responsive to naloxone, contrary to common assumptions made by both medical professionals and in the media (*7*). Both males and females exhibited some level of somatic withdrawal behaviors to xylazine and naloxone, though females showed enhanced behavioral responding. In fact, females appear to be as sensitive, if not more sensitive to xylazine than fentanyl (at the doses tested), while males remain much more responsive to fentanyl conditions. These interesting findings led us to consider the possibility of direct xylazine activity on opioid receptors. Previous studies have shown that xylazine can result in antinociception, results in a cross-tolerance to some mechanisms of opioid induced antinociception, and that these effects are naloxone-sensitive, but surprisingly not sensitive to the κOR selective antagonist nor-binaltorphimine (NorBNI) (*56–59*). Until now, xylazine was thought to exert these effects through promotion of endogenous opioid release and has not been directly tested as a potential opioid agonist. We are the first to report definitive evidence that xylazine acts as a full agonist at κOR.

The interaction of xylazine and fentanyl is intriguing also, because we have previously shown that morphine withdrawal desensitizes α_2_-ARs in outbred rats (Sprague-Dawley) to the level of rats that are genetically predisposed to self-administer opioids (Lewis) (*48*). Further, morphine alone can potentiate norepinephrine (NE) release in the bed nucleus of the stria terminalis (BNST) of the Lewis rats, suggesting complex interactions of opioids on these circuits. As NE in the ventral noradrenergic bundle is critical for opioid reward learning (*60*), the activation of these critical circuits by both opioids and xylazine are targets for future experiments.

These data, along with others (*61, 62*), strongly suggest that there is extensive crosstalk between the α_2_-AR and opioid receptor systems (*48*). Because of this, it is critical to understand how, and if, effects are compounded when agonists target both α_2_-AR and opioid receptors simultaneously. When we tested the ability of an α_2_-AR antagonist (atipamezole) to evoke somatic withdrawal behaviors akin to precipitated opioid withdrawal, we found that again females were more responsive to adrenergic manipulations, even showing a sensitization over days to saline-atipamezole alone. Despite potential differences in responsivity, we found that neither sex was sedated or showed decreased ambulatory activity at the selected dose of 0.5 mg/kg xylazine. Given evidence that in the human population females experience worse withdrawal (*63*), and that female rats self-administer higher levels of fentanyl (*64*), future studies should consider the influence of sex on adrenergic and opioid system interactions.

Both the adrenergic systems and the κOR system are known to have sex differences in rodent models (*65–68*). In our study we found sex differences in locomotion, precipitated withdrawal behavior, and in immediate early gene activation by withdrawal. Similarly, female rats are less sensitive to the depressive effects, and show differential c-Fos activation in the dBNST to κOR agonism when compared with male rats (*69*). Here we also found that withdrawal from fentanyl/xylazine combination increased c-Fos in the dBNST of females but not males. It would be interesting to know which cell types in the dBNST were activated by each of these treatments. In clinical reports, women tend to report enhanced analgesia from mixed κOR/μOR agonists, while rodent models show males with enhanced analgesia to κOR agonism (*70, 71*). These differences may be partially explained by the melanocortin 1 receptor gene and sex differences as related to α-melanocyte-stimulating-hormone (α-MSH) release via κOR dependent mechanisms (*72, 73*). In contrast, it was somewhat surprising to find that c-Fos activity was increased in the LC of both male and female mice in all fentanyl, xylazine, and fentanyl/xylazine combinations in withdrawal (as compared to saline). The LC exhibits sex differences in both form and function with significant receptor and gene expression differences between male and female mice (*66*). Furthermore, μOR agonism is known to differentially affect the discharge rate of LC neurons in male and female rats, likely due to decreased receptor expression (*74*). Future studies will need to examine this circuitry with a more focused lens to determine the role of κOR and the adrenergic systems in mediating the response to fentanyl, xylazine and in combination.

Our findings carry important clinical and public health implications. Considering that xylazine is a full κOR agonist, we note two prominent historical and international examples of non-medical use of the κOR agonist pentazocine: the “Ts and Blues” (pentazocine and tripelennamine) outbreak in the midwestern United States from 1977 to 1981 (*75–77*), and pentazocine injection in Nigeria (*78*) and India (*79–82*). In both settings, characteristic skin lesions beyond the site of injection, eschar formation, and wound cratering were observed (*83–86*), with morphological similarity (*61*) to reports involving xylazine from Puerto Rico (*62*), the Philadelphia area (*16, 63–66*), and New Haven, Connecticut (*67*). κOR distribution in human skin has led to its study as a therapeutic target (*94–96*), suggesting new directions for research into wound etiology. Separately, withdrawal symptoms specific to pentazocine include heightened anxiety, agitation, and paranoia (*71*); these are also cited by clinicians and people who use drugs to be distinguishing presentations of xylazine withdrawal, increasing the difficulty of initiating medication assisted therapy for opioid dependence (*9, 72, 73*). Further investigations are needed to establish if similarities to skin ulcers and withdrawal are coincidental or may be mediated in part by κOR. In addition, existing human pharmaceutical κOR agonists (pentazocine, butorphanol, nalbuphine) could be investigated immediately to alleviate xylazine withdrawal, which is difficult to manage in clinical settings (*7*). Dexmedetomidine, another α_2_-AR agonist that is also approved for human use for other indications in the United States, may also be a candidate medication for investigation to manage xylazine withdrawal.

Current public health and harm reduction messaging makes claims that naloxone is ineffective in reversing the effects of xylazine (*7, 100*). This is problematic because this messaging may lead people to not use naloxone in an overdose scenario when xylazine is suspected in conjunction with fentanyl. In mice, we found that xylazine is responsive to naloxone both in cases where xylazine is administered alone, and in combination with fentanyl. While our findings do not address if xylazine worsens opioid-induced respiratory depression, nor if the presence of xylazine mediates naloxone’s ability to rescue opioid-induced respiratory depression, they do suggest that more nuanced health messaging is warranted in community-based naloxone distribution settings. Opioid-induced respiratory depression is thought to be due to activity at μOR (*101*). Our current data suggest a lack of direct xylazine activity at μOR, however, it is possible that although it has no direct effect on opioid-induced respiratory depression, crosstalk between the two systems or allosteric modulation of μOR might still play a role. The U.S. has seen a recent increase of overdose deaths in which xylazine was identified as contributory to death (*102*); however attribution of causation by medical examiners is inconsistent in practice, and many states do not assay for or report xylazine when present in overdose (*16*). Our findings suggest the urgent need to understand the mechanisms by which xylazine may be implicated in opioid-related overdose, with implications for reporting by forensic medical toxicologists. Our work, and others, seeks to bridge the gap in translatability to continue to provide meaningful animal models of contaminants in the drug supply, providing physicians and regulatory agencies with data to make rapid and effective decisions for public health.

## ACKNOWLEDGMENTS

K_i_ determinations, receptor binding profiles, agonist and/or antagonist functional data, etc. as appropriate was generously provided by the National Institute of Mental Health’s Psychoactive Drug Screening Program, Contract # HHSN-271-2018-00023-C (NIMH PDSP). The NIMH PDSP is Directed by <u>Bryan L. Roth MD</u>, PhD at the University of North Carolina at Chapel Hill and Project Officer <u>Jamie Driscoll</u> at NIMH, Bethesda MD, USA. We thank Maryalice Nocera, Colin Miller, Erin Tracy, Sid Schnoll, and Robyn Jordan for insights into street drug chemical composition and clinical consequences of xylazine use. MLB would like to thank NBB and MRB for their support and help throughout the process of data collection and writing.

## Contributions

Conceptualization: MLB, SYC, SEM, ACN, ND, BLR, ZAM

Validation: MLB, JGM, SEM, SJL, CAC, CGC

Investigation: MLB, JGM, SJL, CAC, CGC, ACN, XPH

Visualization: MLB, JGM, ACN, XPH

Analysis: MLB, JGM, ACN, XPH

Funding acquisition: MLB, BLR, ZAM

Project administration: MLB, ACN, ZAM

Writing – original draft: MLB, JGM, ACN, ZAM

Writing – review & editing: MLB, JGM, SYC ACN, ND, XPH, BLR, ZAM

## SUPPLEMENTAL FIGURES

**Fig. S1.**
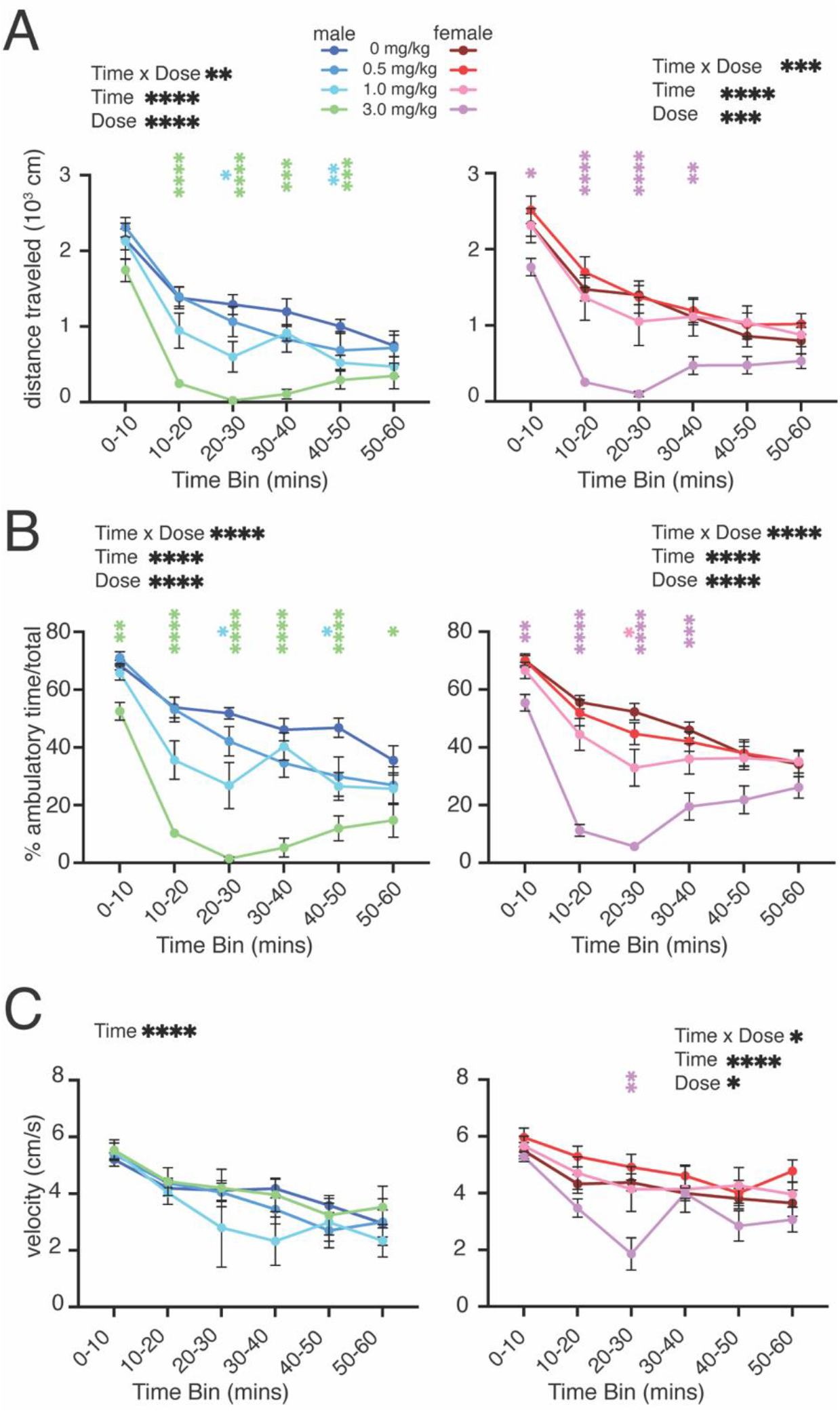
(A) Average distance traveled, (B) % ambulatory time, and (C) velocity of male and female mice administered saline or xylazine (0.5, 1.0, or 3 mg/kg) in 10-minute bins. (A-C) Mixed effects analysis (Time x Dose) with Dunnett’s post-hoc comparisons with saline controls.

**Fig. S2.**
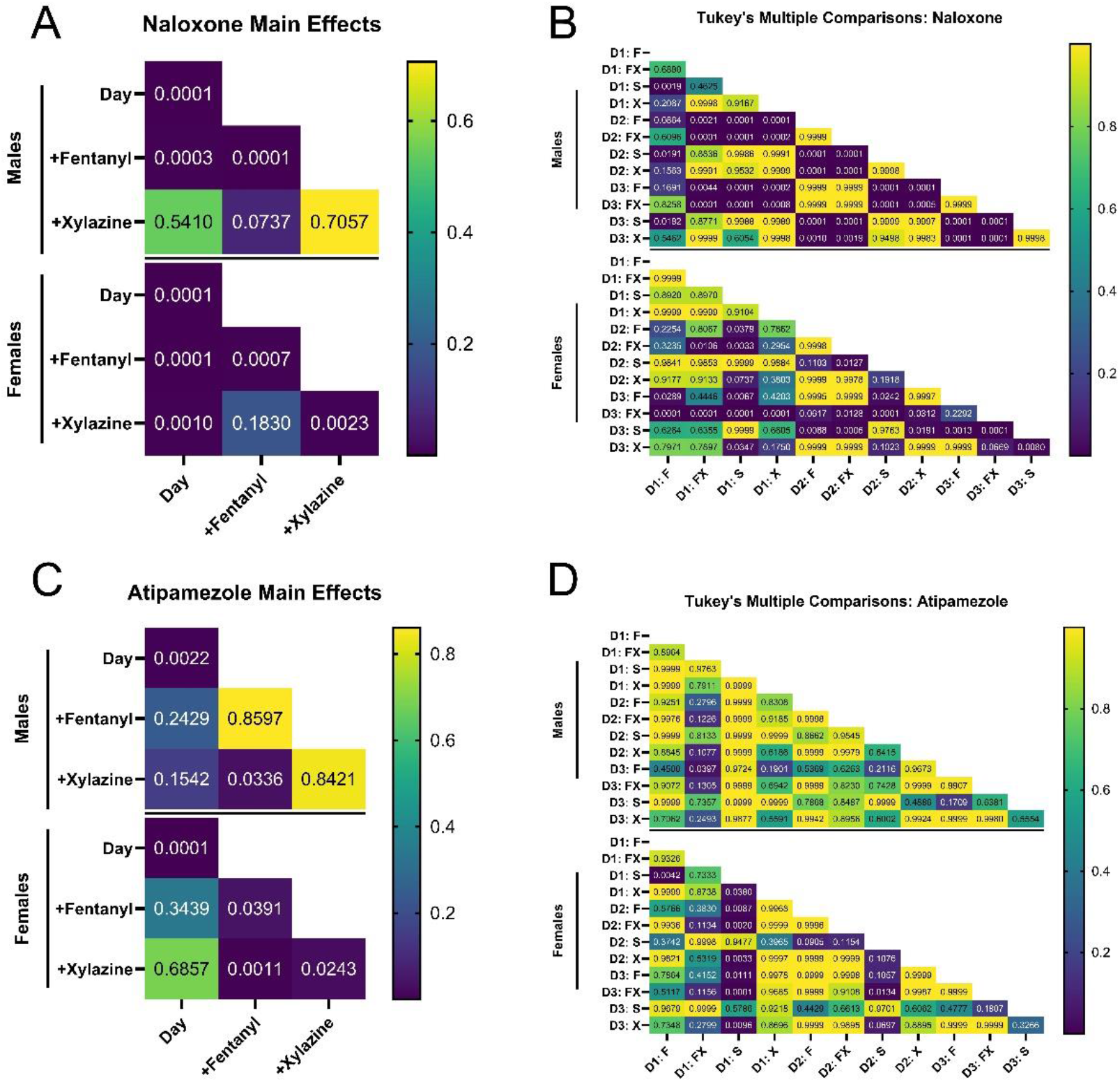
Main effects and Tukey’s post-hoc p-values from precipitated withdrawal. All values shown are p-values. (A,B) Main effects and post hoc from naloxone-precipitated withdrawal corresponding to Fig. 3A,B. (C,D) Main effects and post hoc from atipamezole-precipitated withdrawal corresponding to Fig. 3C,D. Significance p-value=0.05*, 0.1**, 0.001***, <0.0001****.

**Fig. S3.**
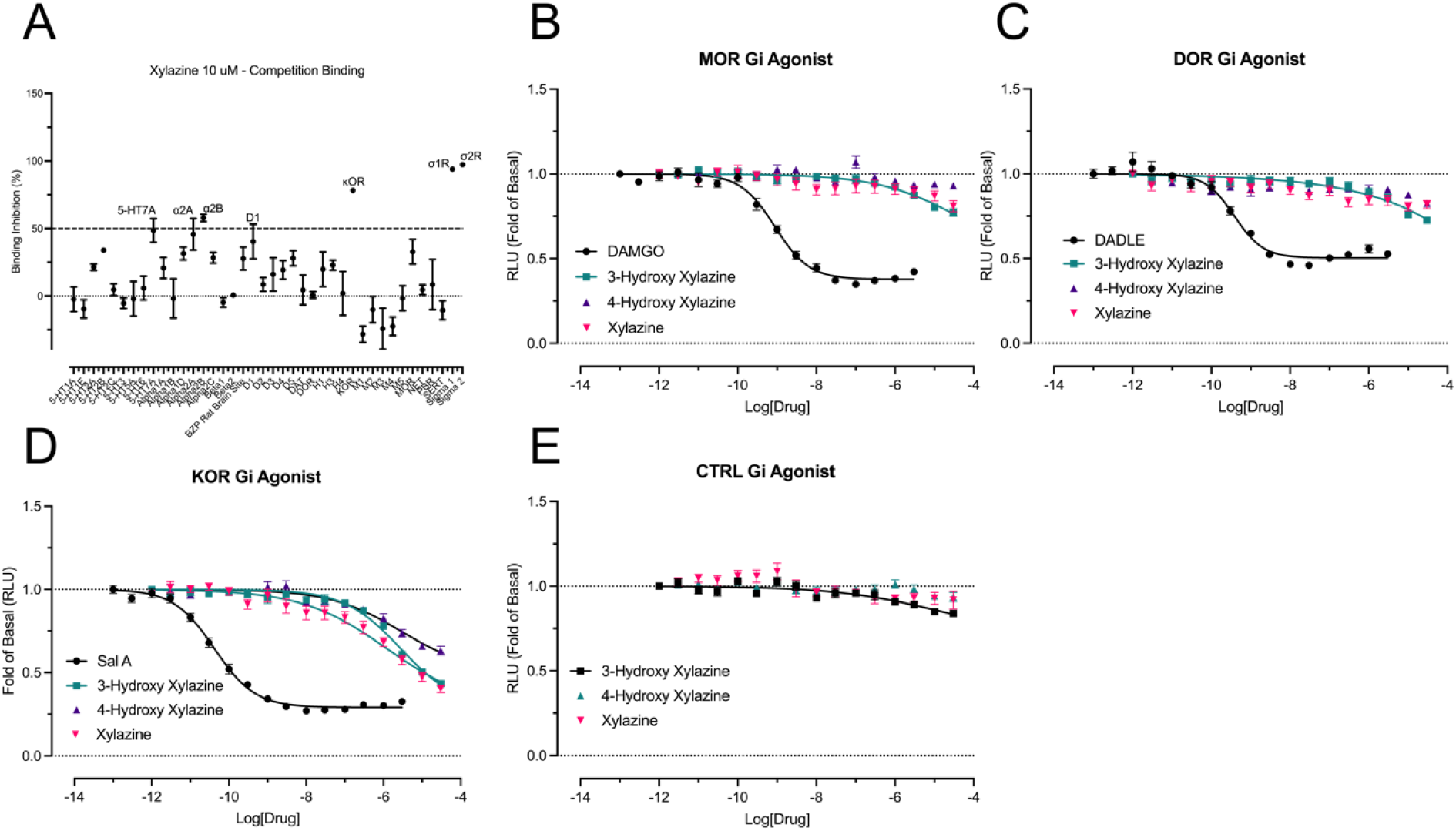
Additional xylazine binding assays. (A) Xylazine binding in radioligand competition binding assay at select receptors. (B-E) Xylazine and metabolite binding at opioid receptors in TRUPATH assay.

## REFERENCES

1. J. I. Elejalde, C. J. Louis, R. Elcuaz, M. A. Pinillos, Drug abuse with inhalated xylazine: European Journal of Emergency Medicine. 10, 252–253 (2003).

2. U. Hoffmann, C. M. Meister, K. Golle, M. Zschiesche, Severe Intoxication with the Veterinary Tranquilizer Xylazine in Humans. Journal of Analytical Toxicology. 25, 245–249 (2001).

3. J. M. Bowles, K. McDonald, N. Maghsoudi, H. Thompson, C. Stefan, D. R. Beriault, S. Delaney, E. Wong, D. Werb, Xylazine detected in unregulated opioids and drug administration equipment in Toronto, Canada: clinical and social implications. Harm Reduct J. 18, 104 (2021).

4. D. G. Spoerke, A. H. Hall, M. J. Grimes, B. N. Honea, B. H. Rumack, Human overdose with the veterinary tranquilizer xylazine. The American Journal of Emergency Medicine. 4, 222–224 (1986).

5. A. G. Gallanosa, D. A. Spyker, J. R. Shipe, D. L. Morris, Human Xylazine Overdose: A Comparative Review with Clonidine, Phenothiazines, and Tricyclic Antidepressants. Clinical Toxicology. 18, 663–678 (1981).

6. R. A. Torruella, Xylazine (veterinary sedative) use in Puerto Rico. Subst Abuse Treat Prev Policy. 6, 7 (2011).

7. R. Gupta, D. R. Holtgrave, M. A. Ashburn, Xylazine — Medical and Public Health Imperatives. N Engl J Med. 388, 2209–2212 (2023).

8. N. Dasgupta, We Can’t Arrest Our Way Out of Overdose: The Drug Bust Paradox. Am J Public Health. 113, 708–708 (2023).

9. F. Montero, P. Bourgois, J. Friedman, Potency-Enhancing Synthetics in the Drug Overdose Epidemic: Xylazine (“Tranq”), Fentanyl, Methamphetamine, and the Displacement of Heroin in Philadelphia and Tijuana. Journal of Illicit Economies and Development. 4, 204–222 (2022).

10. R. Rubin, Here’s What to Know About Xylazine, aka Tranq, the Animal Tranquilizer Increasingly Found in Illicit Fentanyl Samples. JAMA. 329, 1904–1906 (2023).

11. Mackintosh, Potential antidote for Rompun (xylazine) in humans. N Z Med J (1985), doi:3863045.

12. K. L. Rock, A. J. Lawson, J. Duffy, A. Mellor, R. Treble, C. S. Copeland, The first drug-related death associated with xylazine use in the UK and Europe. Journal of Forensic and Legal Medicine. 97, 102542 (2023).

13. R. Ehrman-Dupre, C. Kaigh, M. Salzman, R. Haroz, L.-K. Peterson, R. Schmidt, Management of Xylazine Withdrawal in a Hospitalized Patient: A Case Report. Journal of Addiction Medicine. 16 (2022), doi:10/gsk8ch.

14. S. Ayub, S. Parnia, K. Poddar, A. K. Bachu, A. Sullivan, A. M. Khan, S. Ahmed, L. Jain, Xylazine in the Opioid Epidemic: A Systematic Review of Case Reports and Clinical Implications. Cureus (2023), doi:10/gsnxb8.

15. M. Kariisa, J. O’Donnell, S. Kumar, C. L. Mattson, B. A. Goldberger, Illicitly Manufactured Fentanyl–Involved Overdose Deaths with Detected Xylazine — United States, January 2019–June 2022. MMWR Morb. Mortal. Wkly. Rep. 72, 721–727 (2023).

16. J. Friedman, F. Montero, P. Bourgois, R. Wahbi, D. Dye, D. Goodman-Meza, C. Shover, Xylazine spreads across the US: A growing component of the increasingly synthetic and polysubstance overdose crisis (2023).

17. R. S. Alexander, B. R. Canver, K. L. Sue, K. L. Mo ford, Xylazine and Overdoses: Trends, Concerns, and Recommendations. Am J Public Health. 112, 1212–1216 (2022).

18. R. Rubin, Here’s What to Know About Xylazine, aka Tranq, the Animal Tranquilizer Increasingly Found in Illicit Fentanyl Samples. JAMA. 329, 1904 (2023).

19. Biden-Harris Administration Designates Fentanyl Combined with Xylazine as an Emerging Threat to the United States. White House Press Releases (2023), (available at https://www.whitehouse.gov/ondcp/briefing-room/2023/04/12/biden-harrisadministration-designates-fentanylcombined-with-xylazine-as-an-emerging-threat-to-the-united-states/).

20. S. Browning, D. Lawrence, A. Livingston, B. Morris, INTERACTIONS OF DRUGS ACTIVE AT OPIATE RECEPTORS AND DRUGS ACTIVE AT oc2-RECEPTORS ON VARIOUS TEST SYSTEMS.

21. K. Hohlbaum, B. Bert, S. Dietze, R. Palme, H. Fink, C. Thöne-Reineke, Impact of repeated anesthesia with ketamine and xylazine on the well-being of C57BL/6JRj mice. PLoS ONE. 13, e0203559 (2018).

22. M. Levin-Arama, L. Abraham, T. Waner Harmelin, D. M. Steinberg, T. Lahav, M. Harlev, Subcutaneous Compared with Intraperitoneal Ketamine–Xylazine for Anesthesia of Mice. Journal of the American Association for Laboratory Animal Science. 55 (2016).

23. M. Samini, A. Kardan, S. E. Mehr, Alpha-2 agonists decrease expression of morphine-induced conditioned place preference (2008).

24. T. Uskur, M. A. Barlas, A. G. Akkan, A. Shahzadi, T. Uzbay, Dexmedetomidine induces conditioned place preference in rats: Involvement of opioid receptors. Behavioural Brain Research. 296, 163–168 (2016).

25. S. N. Khatri, S. Pauss, D. Luo, T. E. Prisinzano, K. E. Dunn, J. A. Marusich, J. S. Beckmann, T. D. H. Jr, C. D. Gipson, Xylazine Suppresses Fentanyl Consumption During Self-Administration and Induces a Unique Sex-Specific Withdrawal Syndrome That Is Not Altered By Naloxone in Rats.

26. M. Beuger, A. Tommasello, R. Schwartz, M. Clinton, Clonidine Use and Abuse Among Methadone Program Applicants and Patients. Journal of Substance Abuse Treatment. 15, 589–593 (1998).

27. T. Conway, A. Balson, Concomitant abuse of clonidine and heroin. South Med J. 86, 954–956 (1993).

28. S. J. Dennison, CLONIDINE ABUSE AMONG OPIATE ADDICTS. Psychiatric Quarterly (2001).

29. C. W. Fetrow, J. W. Hoyt, T. M. White, Clonidine patch: reservoir for abuse. J Clin Psychiatry. 55, 266 (1994).

30. P. Lauzon, Two cases of clonidine abuse/dependence in methadone-maintained patients. Journal of Substance Abuse Treatment. 9, 125–127 (1992).

31. J. Schaut, S. H. Schnoll, Four cases of clonidine abuse. Am J Psychiatry. 140, 1625–1627 (1983).

32. E. A. D. Schindler, D. J. Tirado-Morales, D. Kushon, Clonidine Abuse in a Methadone-Maintained, Clonazepam-Abusing Patient. Journal of Addiction Medicine. 7, 218–219 (2013).

33. E. M. Weerts, R. R. Griffiths, Evaluation of the intravenous reinforcing effects of clonidine in baboons. Drug and Alcohol Dependence. 53, 207–214 (1999).

34. W. L. Woolverton, W. D. Wessinger, R. L. Balster, Reinforcing properties of clonidine in rhesus monkeys. Psychopharmacology. 77, 17–23 (1982).

35. I. M. Bravo, B. R. Luster, M. E. Flanigan, P. J. Perez, E. S. Cogan, K. T. Schmidt, Z. A. McElligott, Divergent behavioral responses in protracted opioid withdrawal in male and female C57BL/6J mice. Eur J Neurosci. 51, 742–754 (2020).

36. B. R. Luster, E. S. Cogan, K. T. Schmidt, D. Pati, M. M. Pina, K. Dange, Z. A. McElligott, Inhibitory transmission in the bed nucleus of the stria terminalis in male and female mice following morphine withdrawal. Addiction Biology. 25 (2020), doi:10.1111/adb.12748.

37. M. L. Bedard, J. S. Lord, P. J. Perez, I. M. Bravo, A. T. Teklezghi, L. M. Tarantino, G. H. Diering, Z. A. McElligott, Probing different paradigms of morphine withdrawal on sleep behavior in male and female C57BL/6J mice. Behavioural Brain Research. 448, 114441 (2023).

38. I. Bravo, M. Bluitt, Z. McElligott, “Examining opioid withdrawal scoring and adaptation of global scoring systems to male and female C57BL/6J mice” (preprint, Neuroscience, 2021),, doi:10.1101/2021.10.11.463944.

39. J. Besnard, G. F. Ruda, V. Setola, K. Abecassis, R. M. Rodriguiz, X.-P. Huang, S. Norval, M. F. Sassano, A. I. Shin, L. A. Webster, F. R. C. Simeons, L. Stojanovski, A. Prat, N. G. Seidah, D. B. Constam, G. R. Bickerton, K. D. Read, W. C. Wetsel, I. H. Gilbert, B. L. Roth, A. L. Hopkins, Automated design of ligands to polypharmacological profiles. Nature. 492, 215–220 (2012).

40. W. K. Kroeze, M. F. Sassano, X.-P. Huang, K. Lansu, J. D. McCorvy, P. M. Giguère, N. Sciaky, B. L. Roth, PRESTO-Tango as an open-source resource for interrogation of the druggable human GPCRome. Nat Struct Mol Biol. 22, 362–369 (2015).

41. L. Salas-Estrada, D. Provasi, X. Qiu, H. Ü. Kaniskan, X.-P. Huang, J. F. DiBerto, J. M. Lamim Ribeiro, J. Jin, B. L. Roth, M. Filizola, De Novo Design of κ-Opioid Receptor Antagonists Using a Generative Deep-Learning Framework. J. Chem. Inf. Model. 63, 5056–5065 (2023).

42. R. H. J. Olsen, J. F. DiBerto, J. G. English, M. Gl udin, B. E. Krumm, S. T. Slocum, T. Che, A. C. Gavin, J. D. McCorvy, B. L. Roth, R. T. Strachan, TRUPATH, an open-source biosensor platform for interrogating the GPCR transducerome. Nat Chem Biol. 16, 841–849 (2020).

43. T. Kitano, T. Kobayashi, S. Yamaguchi, K. Otsuguro, The α 2A -adrenoceptor subtype plays a key role in the analgesic and sedative effects of xylazine. J vet Pharmacol Therap. 42, 243–247 (2019).

44. M. Samini, A. Kardan, S. Mehr, Alpha-2 agonists decrease expression of morphine-induced conditioned place preference. Pharmacology Biochemistry and Behavior. 88, 403–406 (2008).

45. G. F. Koob, M. Le Moal, Drug Abuse: Hedonic Homeostatic Dysregulation. Science. 278, 52–58 (1997).

46. G. F. Koob, M. Le Moal, Drug Addiction, Dysregulation of Reward, and Allostasis. 24, 34 (2001).

47. G. F. Koob, N. D. Volkow, Neurobiology of addiction: a neurocircuitry analysis. The Lancet Psychiatry. 3, 760–773 (2016).

48. Z. A. McElligott, M. E. Fox, P. L. Walsh, D. J. Urb n, M. S. Ferrel, B. L. Roth, R. M. ightman, Noradrenergic Synaptic Function in the Bed Nucleus of the Stria Terminalis Varies in Animal Models of Anxiety and Addiction. Neuropsychopharmacol. 38, 1665–1673 (2013).

49. G. Schulteis, C. J. Heyser, G. F. Koob, Differential expression of response-disruptive and somatic indices of opiate withdrawal during the initiation and development of opiate dependence: Behavioural Pharmacology. 10, 235–242 (1999).

50. E. M. Lefevre, M. T. Pisansky, C. Toddes, F. Baruffaldi, M. Pravetoni, L. Tian, T. J. Y. Kono, P. E. Rothwell, Interruption of continuous opioid exposure exacerbates drug-evoked adaptations in the mesolimbic dopamine system. Neuropsychopharmacol. 45, 1781–1792 (2020).

51. E. M. Lefevre, E. A. Gauthier, L. L. Bystrom, J. Scheunemann, P. E. Rothwell, J. Neurosci., in press, doi:10.1523/JNEUROSCI.0595-22.2022.

52. L. Chung, A Brief Introduction to the Transduction of Neural Activity into Fos Signal. Development & Reproduction. 19, 61–67 (2015).

53. A. Chaudhuri, S. Zangenehpour, F. Ye, Molecular maps of neural activity and quiescence.

54. D. S.-G. Lavoie, F. Pailleux, P. Vachon, F. Beaudry, Characterization of xylazine metabolism in rat liver microsomes using liquid chromatography-hybrid triple quadrupole-linear ion trap-mass spectrometry: Characterization of xylazine metabolism. Biomed. Chromatogr. 27, 882–888 (2013).

55. M.-H. Spyridaki, E. Lyris, I. Georgoulakis, D. Kouretas, M. Konstantinidou, C. G. Georgakopoulos, Determination of xylazine and its metabolites by GC–MS in equine urine for doping analysis. Journal of Pharmaceutical and Biomedical Analysis. 35, 107–116 (2004).

56. T. D. Ware, D. Paul, Cross-tolerance between analgesia produced by xylazine and selective opioid receptor subtype treatments. European Journal of Pharmacology. 389, 181–185 (2000).

57. T. R. L. Romero, D. D. F. Pacheco, I. D. G. Duarte, Xylazine induced central antinociception mediated by endogenous opioids and μ-opioid receptor, but not δ-or κ-opioid receptors. Brain Research. 1506, 58–63 (2013).

58. T. R. L. Romero, A. De Castro Perez, J. N. De Francischi, I. D. Gama Duarte, Probable involvement of α2Cadrenoceptor subtype and endogenous opioid peptides in the peripheral antinociceptive effect induced by xylazine. European Journal of Pharmacology. 608, 23–27 (2009).

59. M. Nakamura, S. H. Ferreira, Peripheral analgesic action of clonidine: mediation by release of endogenous enkephalin-like substances. European Journal of Pharmacology. 146, 223–228 (1988).

60. V. G. Olson, C. L. Heusner, R. J. Bland, M. . During, D. Weinshenker, R. D. Palmiter, Role of Noradrenergic Signaling by the Nucleus Tractus Solitarius in Mediating Opiate Reward. Science. 311, 1017–1020 (2006).

61. G. K. Aghajanian, Tolerance of locus coeruleus neurones to morphine and suppression of withdrawal response by clonidine. Nature. 276, 186–188 (1978).

62. G. K. Aghajanian, J. H. Kogan, B. Moghaddam, Opiate withdrawal increases glutamate and aspartate efflux in the locus coeruleus: an in vivo microdialysis study. Brain Research. 636, 126–130 (1994).

63. O. D. Ware, J. D. Ellis, K. E. Dunn, J. G. Hobelmann, P. Finan, A. S. Huhn, The association of chronic pain and opioid withdrawal in men and women with opioid use disorder. Drug and Alcohol Dependence. 240, 109631 (2022).

64. E. B. Towers, B. Setaro, W. J. Lynch, Sex- and Dose-Dependent Differences in the Development of an Addiction-Like Phenotype Following Extended-Access Fentanyl Self-Administration. Front. Pharmacol. 13, 841873 (2022).

65. P. Sun, J. Wang, M. Zhang, X. Duan, Y. Wei, F. Xu, Y. Ma, Y.-H. Zhang, Sex-Related Differential Whole-Brain Input Atlas of Locus Coeruleus Noradrenaline Neurons. Front. Neural Circuits. 14, 53 (2020).

66. B. Mulvey, D. L. Bhatti, S. Gyawali, A. M. Lake, S. Kriaucionis, C. P. Ford, M. R. Bruchas, N. Heintz, J. D. Dougherty, Molecular and Functional Sex Differences of Noradrenergic Neurons in the Mouse Locus Coeruleus. Cell Reports. 23, 2225–2235 (2018).

67. E. H. Chartoff, M. Mavrikaki, Sex Differences in Kappa Opioid Receptor Function and Their Potential Impact on Addiction. Front. Neurosci. 9 (2015), doi:10/gccxgz.

68. D. A. Bangasser, K. R. Wiersielis, S. Khantsis, Sex differences in the locus coeruleus-norepinephrine system and its regulation by stress. Brain Research. 1641, 177–188 (2016).

69. S. E. Russell, A. B. Rachlin, K. L. Smith, J. Muschamp, L. Berry, Z. Zhao, E. H. Chartoff, Sex Differences in Sensitivity to the Depressive-like Effects of the Kappa Opioid Receptor Agonist U-50488 in Rats. Biological Psychiatry. 76, 213–222 (2014).

70. R. W. Gear, N. C. Gordon, P. H. Heller, S. Paul, C. Miaskowski, J. D. Levine, Gender difference in analgesic response to the kappa-opioid pentazocine. Neuroscience Letters. 205, 207–209 (1996).

71. A. C. Barrett, C. D. Cook, J. M. Terner, E. L. Roach, C. Syvanthong, M. J. Picker, Sex and rat strain determine sensitivity to κ opioid-induced antinociception. Psychopharmacology. 160, 170–181 (2002).

72. J. S. Mogil, S. G. Wilson, E. J. Chesler, A. ankin, K. V. S. Nemmani, W. R. Lariviere, M. K. Groce, M. R. Wallace, L. Kaplan, R. Staud, T. J. Ness, T. L. Glover ankova, A. Mayorov, V. J. Hruby, J. E. Grisel, R. B. Fillingim, The melanocortin-1 receptor gene mediates female-specific mechanisms of analgesia in mice and humans. Proc. Natl. Acad. Sci. U.S.A. 100, 4867–4872 (2003).

73. J. Manzanares, E. J. Wagner, K. E. Moore, K. J. Lookingland, Kappa opioid receptormediated regulation of prolactin and amelanocyte-stimulating hormone secretion in male and female rats. Life Sciences. 53, 795–801 (1993).

74. H. M. Guajardo, K. Snyder, A. Ho, R. J. Valentino, Sex Differences in μ-Opioid Receptor Regulation of the Rat Locus Coeruleus and Their Cognitive Consequences. Neuropsychopharmacol. 42, 1295–1304 (2017).

75. A. Poklis, Pentazocine/tripelennamine (T’s and Blues) abuse: A five year survey of St. Louis, Missouri. Drug and Alcohol Dependence. 10, 257–267 (1982).

76. G. B. Irons, D. J. Hodgkinson, G. C. Chong, J. E. Woods, Pentazocine Ulceration. 2, 286–289 (1970).

77. R. S. Padilla, L. E. Becker, H. Hoffman, G. Long, Cutaneous and Venous Complications of Pentazocine Abuse. Archives of Dermatology. 115, 975–977 (1979).

78. C. Otene, I. Ohiaeri, D. Odatuwa-Omagbemi, R. E. T. Enemudo, Abuse of parenteral opioid (pentazocine) amongst plastic surgery patients in a tertiary health institution in southsouth Nigeria − a case series. Nigerian Journal of Plastic Surgery. 16 (2020).

79. A. Pawar, A. K. Rajalakshmi, R. P. Upadhyay, Pentazocine use among people who inject drugs in India. Asian Journal of Psychiatry. 16, 3–6 (2015).

80. S. Sethi, R. Sarkar, V. Garg, N. Khurana, Pentazocine-induced ulcers revisited. International Journal of Dermatology. 55, e49–e51 (2016).

81. V. M. Agashe, H. Patil, M. K. Gundavda, Multiple Skin Abscesses and Myofibrosis of Bilateral Lower Limbs Following Repeated Intramuscular Injection of Pentazocine with Concomitant Tuberculous Infection. Journal of Orthopaedic Case Reports. 5 (2015).

82. S. Yelne, M. B. Wanjari, P. K. Munjewar, K. Malu, R. Umate, A Rare Incidence of Necrotizing Lesion of the Upper Arm Due to Pentazocine Injection Abuse: A Case Report. Cureus (2022), doi:10/gsk7s2.

83. S. Kumar, S. Chaudhury, S. Soren, J. Simlai, R. Kumari, Local complications of pentazocine abuse: Case report and review. Ind Psychiatry J. 27, 296 (2018).

84. J. E. Schlicher, R. L. Zuehlke, P. J. Lynch, Local Changes at the Site of Pentazocine Injection. Archives of Dermatology. 104, 90–91 (1971).

85. D. L. Parks, H. O. Perry, S. A. Muller, Cutaneous Complications of Pentazocine Injections. Archives of Dermatology. 104, 231–235 (1971).

86. R. F. Palestine, J. L. Millns, G. T. Spigel, L. Schroeter, Skin manifestations of pentazocine abuse. Journal of the American Academy of Dermatology. 2, 47–55 (1980).

87. A. Bishnoi, V. Singh, U. Khanna, K. Vinay, Skin ulcerations caused by xylazine: A lesser-known entity. Journal of the American Academy of Dermatology. 89, e99–e102 (2023).

88. J. C. Reyes, J. L. Negrón, H. M. Colón, A. M. P illa, M. Y. Millán, T. D. Matos, R. R. Robles, The Emerging of Xylazine as a New Drug of Abuse and its Health Consequences among Drug Users in Puerto Rico. J Urban Health. 89, 519–526 (2012).

89. S. V. Malayala, B. N. Papudesi, R. Bobb, W mbush, Xylazine-Induced Skin Ulcers in a Person Who Injects Drugs in Philadelphia, Pennsylvania, USA. Cureus (2022), doi:10/gr9cbw.

90. L. Rose, R. Kirven, K. Tyler, C. Chung, A. M. K man, Xylazine-induced acute skin necrosis in two patients who inject fentanyl. JAAD Case Reports. 36, 113–115 (2023).

91. J. Wei, C. Wachuku, J. Berk-Krauss, K. T. Steele, M. Rosenbach, E. Messenger, Severe cutaneous ulcerations secondary to xylazine (tranq): A case series. JAAD Case Reports. 36, 89–91 (2023).

92. S. Rengifo, A. M. Ilyas, R. Tosti, Upper Extremity Soft Tissue Wound Related to Xylazine-laced Fentanyl Intravenous (IV) Drug Abuse: A Case Report. SurgiColl. 1 (2023), doi:10/gsk8cd.

93. A. Dowton, M. Doernberg, E. Heiman, P. Barelli, M. Golden, H. Wang, J. Leventhal, K. L. Mo ford, K. L. Sue, Recognition and Treatment of Wounds in Persons Using Xylazine: A Case Report From New Haven, Connecticut. Journal of Addiction Medicine (2023) (available at https://journals.lww.com/journaladdiction medicine/fulltext/9900/recognition_and_tr eatment_of_wounds_in_persons.203.aspx) .

94. S. Salemi, A. Aeschlimann, N. Reisch, A. Jüngel, R. E. Gay, F. L. Heppner, B. A. Michel, S. Gay, H. Sprott, Detection of kappa and delta opioid receptors in skin— Outside the nervous system. Biochemical and Biophysical Research Communications. 338, 1012–1017 (2005).

95. B. Cheng, H. Liu, X. Fu, Z. Sheng, J. Li, Coexistence and upregulation of three types of opioid receptors, mu, delta and kappa, in human hypertrophic scars. British Journal of Dermatology. 158, 713–720 (2008).

96. L. M. Snyder, M. C. Chiang, E. Loeza-Alcocer, Y. Omori, J. Hachisuka, T. D. Sheahan, J. R. Gale, P. C. Adelman, E. I. Sypek, S. A. Fulton, R. L. Friedman, M. C. Wright, M. G. Duque, Y. S. Lee, Z. Hu, H. Huang, X. Cai, K. A. Meerschaert, V. Nagarajan, T. Hirai, G. Scherrer, D. H. Kaplan, F. Porreca, B. M. Davis, M. S. Gold, H. R. Koerber, S. E. Ross, Kappa Opioid Receptor Distribution and Function in Primary Afferents. Neuron. 99, 1274–1288.e6 (2018).

97. W. R. Lange, D. R. Jasinski, The clinical pharmacology of pentazocine and tripelennamine (T’s and Blues). Advances in Alcohol & Substance Abuse. 5, 71–83 (1986).

98. T. Boyce, G. D. Whitehill, B. J. Anderson, “Management of Xylazine Withdrawal in a Patient Admitted to the Intensive Care Unit With Carbon Monoxide Poisoning” in C48. CASE REPORTS: TOXICOLOGY AND OTHER INVOLVING MEDICATIONS (American Thoracic Society, 2023; https://www.atsjournals.org/doi/10.1164/ajrccm-conference.2023.207.1_MeetingAbstracts.A5204), pp. A5204–A5204.

99. A. Spadaro, A. Spadaro, K. O’Connor, S. Lakamana, A. Sarker, R. Wightman, Self-reported Xylazine Experiences: A Mixed Methods Study of Reddit Subscribers.

100. N. Chhabra, M. Mir, M. J. Hua, S. Berg, J. Nowinski-Konchak, S. Aks, P. Arunkumar, K. Hinami, Notes From the Field: Xylazine-Related Deaths — Cook County, Illinois, 2017–2021. MMWR Morb. Mortal. Wkly. Rep. 71, 503–504 (2022).

101. M. Z. Imam, A. Kuo, S. Ghassabian, M. T. Smith, Progress in understanding mechanisms of opioid-induced gastrointestinal adverse effects and respiratory depression. Neuropharmacology. 131, 238–255 (2018).

102. M. Spencer, J. Cisewski, M. Warner, M. Garnett, “Drug Overdose Deaths Involving Xylazine, United States, 2018–2021” (National Center for Health Statistics (U.S.), 2023),, doi:10.15620/cdc:129519.

